# Laterally biased diffusion of males of the water flea *Daphnia magna*

**DOI:** 10.1101/2021.01.21.427564

**Authors:** Kenji Toyota, Masaki Yasugi, Norihisa Tatarazako, Taisen Iguchi, Eiji Watanabe

## Abstract

The water flea *Daphnia magna* is a representative example of zooplankton living in freshwater environments. They primarily propagate via asexual reproduction under normal and healthy environmental conditions. Environmental stimuli that signal a shift to disadvantageous conditions induce *D. magna* to change their mode of reproduction from asexual to sexual reproduction. During the sexual reproduction phase, they produce special tough eggs (resting eggs), which can survive severe environmental conditions. Despite our increased understanding of their mating behaviours, the sex-specific characteristics of swimming behaviours among daphnid species are poorly understood. In this study, we analysed the swimming patterns and dynamics of female and male adult *D. magna*. First, we found laterally biased diffusion of males in contrast to the homogeneous, nondirectional diffusion of females. Second, computer modelling analysis using a discrete-time Markov chain simulation, in which the frequencies of turning behaviour were evaluated as probability distributions, explained the greater diffusion of males in the horizontal direction. Under the presumption that high diffusion in the horizontal direction increases the probability of encountering a distant mate, these findings led us to hypothesise that male *D. magna* increase genotype heterogeneity by effectively selecting the probability distributions of certain motion parameters.

**Summary statements:** We analysed the swimming behaviours of adult water flea *Daphnia magna*, and found apparent sexual differences: laterally biased diffusion of males in contrast to the nondirectional diffusion of females.

## Introduction

For animals, survival depends on the ability to engage in appropriate behaviours in response to current circumstances. In various situations, animals must perform a range of behaviours, such as feeding, escape, aggression and sexual behaviours, by appropriately regulating their motor systems. Daphnids (referred to as water fleas), which are cladoceran crustaceans living in freshwater ecosystems, are no exception. Although they cannot perform smooth motion in the same manner as higher-order animals, such as humans, they can engage in appropriate behaviours to search for food, escape from predators and find mates using primitive sensors and motors (Ebert, 2005). In this study, we studied the swimming behaviours of *Daphnia magna* as a representative example of daphnid species (Figure 1). It is one of the most readily available and large species of daphnids, and it occupies a key position in food webs of shallow ponds.

**Figure 1.**
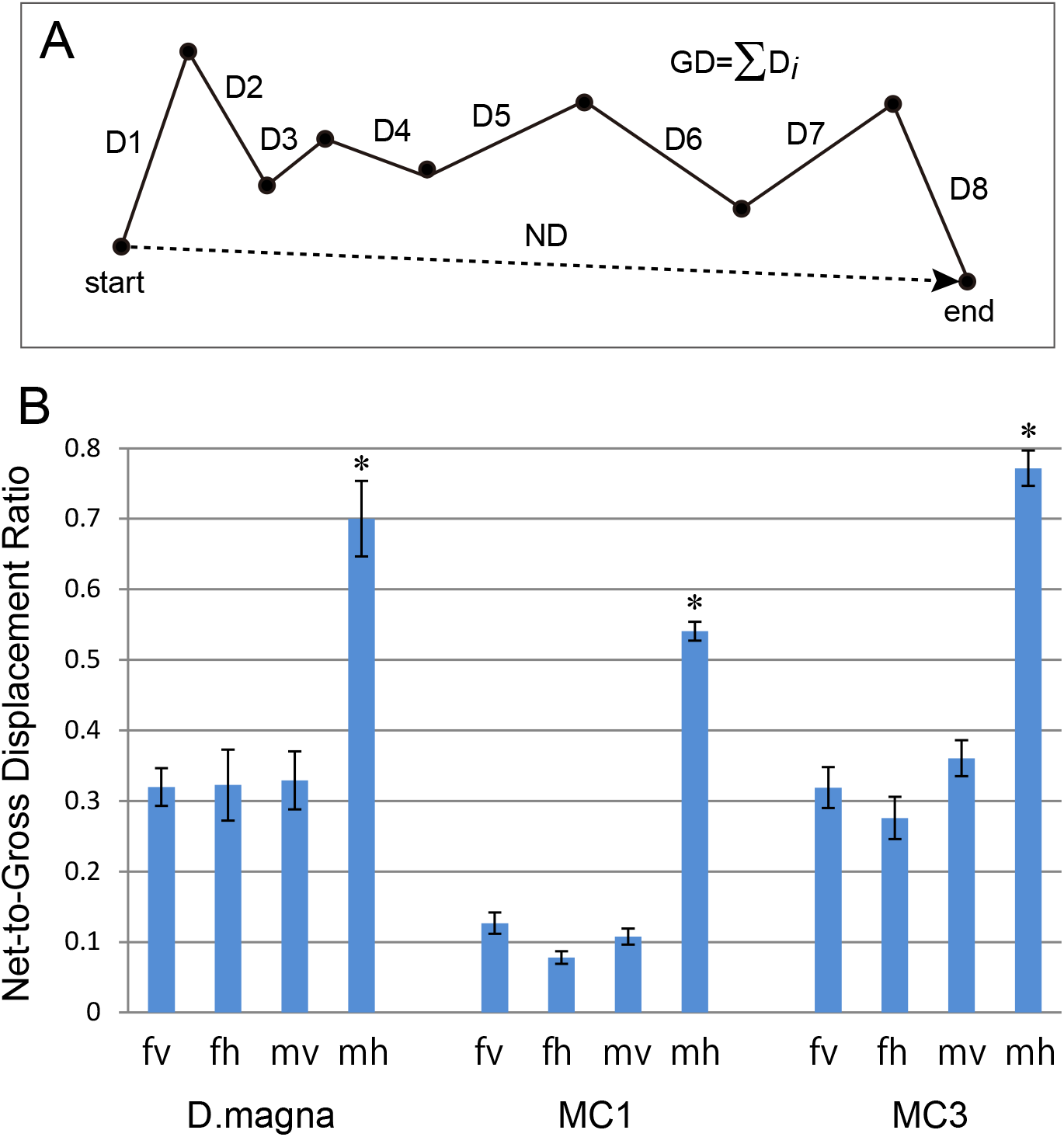
*D. magna*. Sexually mature female (left) and male (right) *D. magna* (lateral view). Scale bar, 1 mm.

The jumping-like motion of daphnids stems from the beating of their second antennae, which do not act as sensory organs but rather function in a manner similar to the oars of a boat (refer to supplemental movies 1 and 2). Their rapid downbeat produces a swift upward movement, whereas motionless organisms rapidly sink due to gravity (Haury and Weihs, 1976; Matsunaga and Watanabe, 2012). This rhythmic swimming pattern results in directed movement. In addition, these organisms occasionally drastically change their direction via irregular and disordered motion of their second antennae (Garcia et al., 2007). By simply repeating downbeats, succumbing to the pull of gravity and occasionally performing turning behaviour, they perform vertical and horizontal migration towards or away from a specific location in a pond or a lake. Such swimming patterns in a random fashion are widely observed in zooplankton species (Seuront and Strutton, 2003). Although these organisms freely move in water under constant conditions in the absence of specific physical stimuli, such as water flow or directional light, the motion of zooplankton is fundamentally explained by random walks or Levy flights (Bartumeus et al., 2003; Garcia et al., 2007; Kiørboe and Bagøien, 2005; Seuront et al., 2004a; 2004b; Visser and Thygesen, 2003), which are representative types of Markov chain (MC) processes. The movement vector (motion direction and movement length) at each step is stochastically determined. In addition to the random walk theory, mathematical models based on continuous stochastic differential equations closely approximate zooplankton behaviours (Grünbaum, 2000; Øien, 2004; Schimansky-Geier et al., 2005). Biologists might expect that these internal stochastic processes found in zooplankton play an important role in their biological behaviours, such as foraging (Bartumeus et al., 2003; Garcia et al., 2007).

Among the variety of characterised behaviours of cladocera species, including daphnids, mating behaviours, such as encounters, grasping and copulation, have received growing attention (Brewer, 1998; Kerfoot, 1980; Winsor and Innes, 2002), given that daphnids have evolved a unique reproductive system for increasing the encounter probability between female and male individuals in a population. Their method of reproduction is termed cyclical parthenogenesis, in which parthenogenetic (asexual) and sexual phases are interchanged in response to external environmental stimuli, such as day length, temperature, nutrient availability and crowding (Banta and Brawn, 1929; Hobæk and Larsson, 1990; Kleiven et al., 1992; Smith, 1915). Under favourable growth conditions, they parthenogenetically produce diploid eggs, giving rise to genetically identical female offspring. When environmental conditions deteriorate, they produce males and sexual females bearing haploid diapause eggs, permitting sexual reproduction and increasing genetic diversity. Resting eggs are resistant to harsh conditions, such as drying, freezing and exposure to the digestive enzymes of fish (Jarnagin et al., 2000). Despite this accumulated knowledge on daphnid mating behaviours, little information concerning sexual differences in swimming behaviours among daphnid species is available (Brewer, 1998). The goal of the present study was to elucidate the sex-specific swimming patterns of adult daphnids and to reproduce these observed behavioural patterns via computer modelling. We found laterally biased diffusion of males in contrast to the nondirectional diffusion of females. Computer simulation showed that the frequencies of turning behaviour are involved in male-specific behaviour.

## Results

### Overview of swimming patterns

Figures 2A and 2B show the swimming trajectories of 28 female and 28 male *D. magna* individuals. They travelled on average 111.52 ± 2.42 mm (females) and 96.17 ± 2.68 mm (males) per individual from their starting position to their ending position in 8.53 s (256 frames), with mean speeds of 13.07 ± 0.028 mm/s and 11.27 ± 0.31 mm/s, respectively. In Figures 2A and 2B, consecutive coordinates were normalised to the given halfway point (the 129^th^ coordinate). To reflect all data of the coordinates of the trajectory in a single graph, the trajectory graph was transformed into a probability density plot (Figures 2C and 2D). Next, we performed kernel density estimation. Whereas females exhibited a relatively symmetric distribution of movement in all directions, males exhibited a biased movement distribution in the horizontal direction. The horizontal-to-vertical ratio of high-density areas (density of 0.6 and greater; greenish yellow to white in Figures 2C and 2D) was 1.35 in females and 2.10 in males. The highest density for females was 1.16-fold greater than that for males. Furthermore, a comparison of the area with a density of at least the highest density for males (density of 1.1 and greater) revealed that the area for females was 4.29-fold larger than that for males.

**Figure 2.**
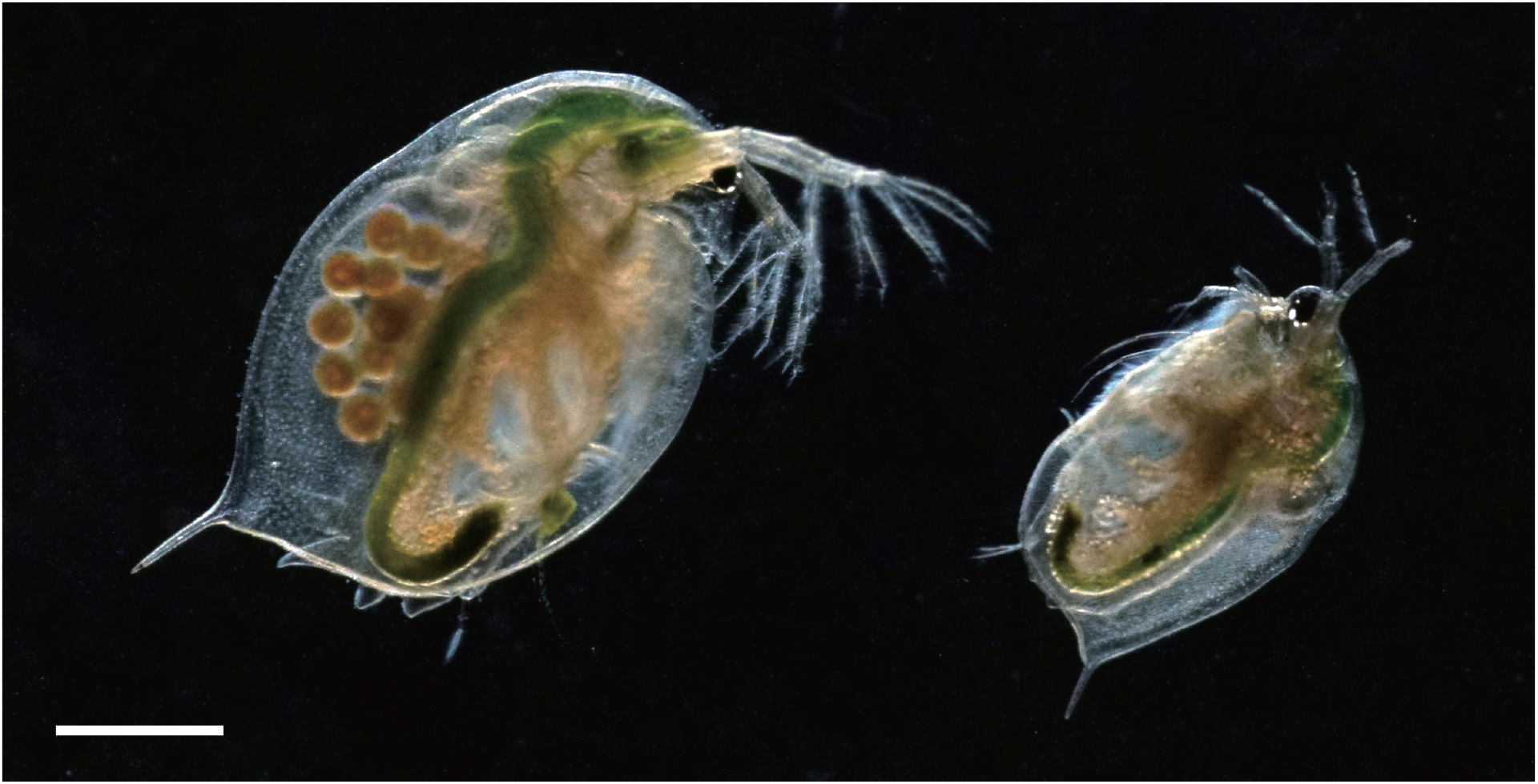
Overview of swimming trajectories. Time series plot of female (A) and male (B) *D. magna*. In total, 7,196 coordinates (240 sets, 257 consecutive coordinates per individual, *n* = 28) of the calculated centre of mass of *D. magna* were plotted along the vertical and horizontal axes. Consecutive coordinates were normalised to the given halfway point (the 129^th^ coordinate), which was set as vertical coordinate 50 and horizontal coordinate 50. Each plot was annotated by arbitrary symbols with different colours (blue, red, green, orange, violet or yellow). Trajectory graphs are represented as probability density plots for females (C) and males (D). Kernel density estimation was performed using a quartic kernel function with a bandwidth of 12 mm. The two-dimensional density map was visualised using terra colours. Scale bar, 10 mm.

### Mean squared displacement and autocorrelation function

Next, we analysed the MSD of the motion of *D. magna.* Figure 3 shows the MSD derived from the trajectories of *D. magna* females and males. The MSD was calculated independently in vertical and horizontal directions, and best-fit lines were calculated using a power-law approximation. The power-law exponent in the vertical direction was 1.58 for females (least mean square = 0.97) and 1.39 for males (0.97). The power-law exponent in the horizontal direction was 1.29 for females (0.93) and 1.82 for males (0.99). The male-to-female ratio of the power-law exponent was 0.88 in the vertical direction and 1.41 in the horizontal direction. The MSD plot of females moving in horizontal directions appears to consist of two time scales divided at approximately 5,000 ms (Figure 3, bottom, red). However, to compare diffusion rates of *D. magna* females and males, each of them was simply represented by a single power-law exponent. All power-law exponents larger than 1.0, indicating diffusion of both *D. magna* females and males, were labelled as ‘superdiffusion’ in both directions. The motion analysis data also indicated that horizontal diffusion rates of males were higher than those of females irrespective of the lower mean speed of males.

**Figure 3.**
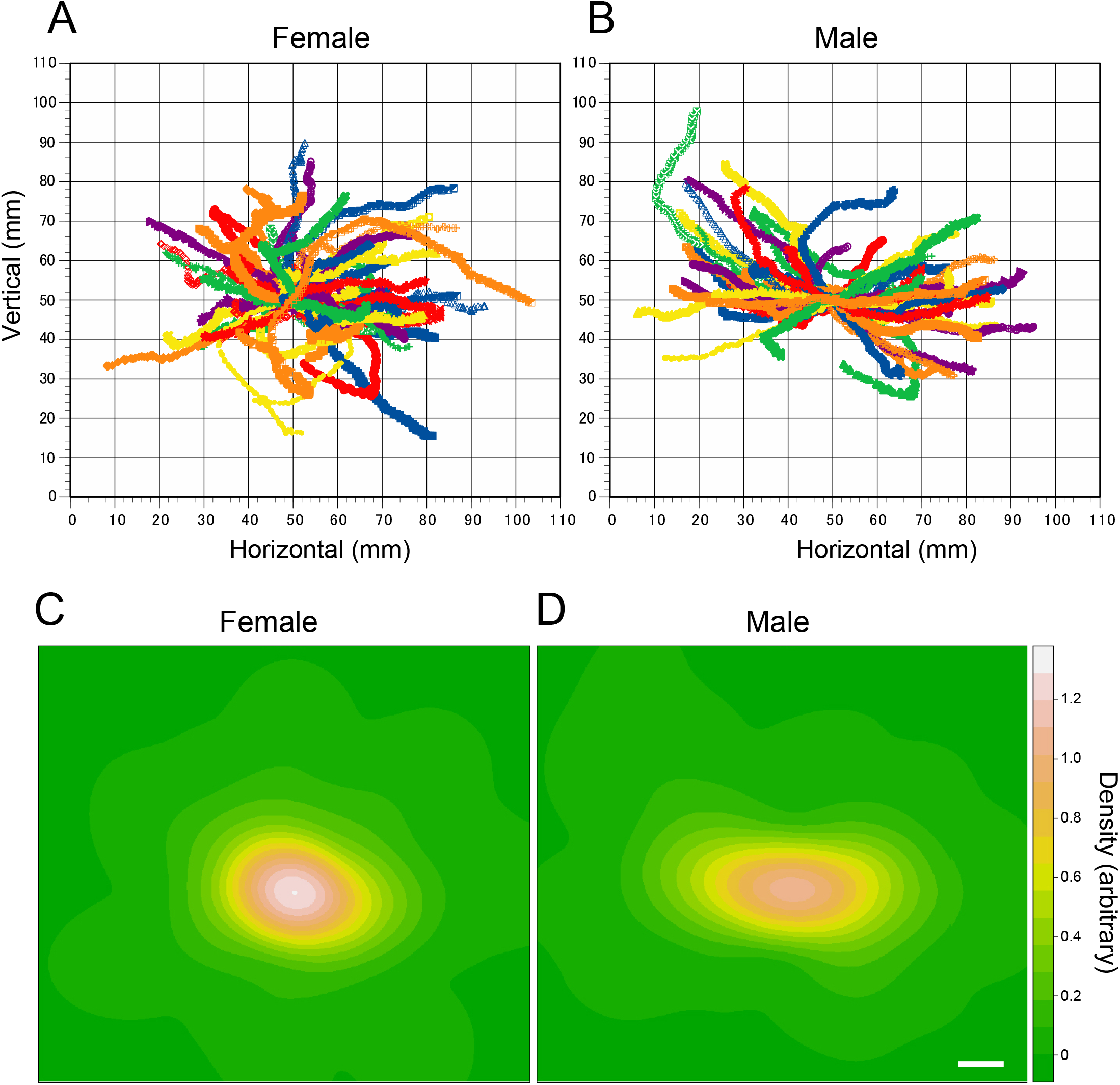
Mean squared displacement of *D. magna*. The mean squared displacements (MSDs) of female (red) and male (blue) *D. magna* are plotted along the vertical (top) and horizontal (bottom) axes (logarithmic scales, *n* = 28). Best-fit lines using a power-law approximation are shown in black. The power-law exponent (α) and least mean square (*R*^2^) are shown in the plots.

The estimate for the correlation timescale obtained from the velocity autocorrelation function is shown in Figure 4. In both cases (for females and males as well as for vertical and horizontal diffusions), distinct periodicity and autocorrelation characteristics were not observed. In many cases, a persistent random walk (or correlated random walk) was used for modelling the trajectories of the moving animals, given that the velocities of moving animals are persistent, in general (Codling et al., 2008). However, the present models include a fundamentally uncorrelated random walk because of the low autocorrelation characteristic. Using statistical MC models based on the discrete probability distributions of the velocities, move-step lengths, and SCISs (turns), we attempted to identify the parameter element contributing to the higher diffusion rate of *D. magna* males in the horizontal direction. The MC3 model referencing the discrete probability distribution of SCIS is a type of persistent random walk, given that the size values of SCIS are coupled with the persistence of the movement direction.

**Figure 4.**
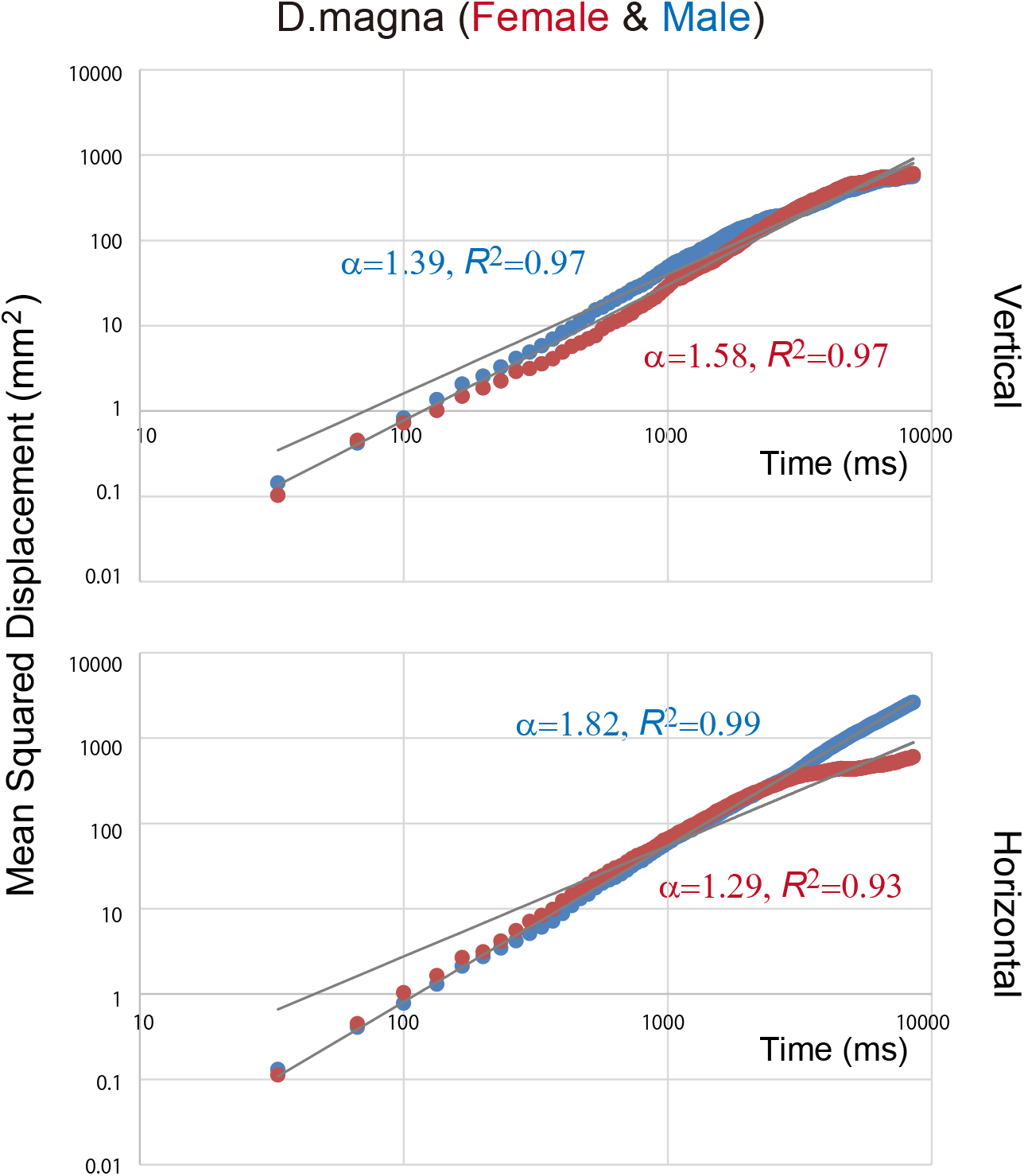
Autocorrelation function of *D. magna*. Average normalised autocorrelation functions of female (top) and male (bottom) *D. magna* are plotted along the vertical (left) and horizontal (right) axes (*n* = 28).

### MC1

Assuming the MC in reference to frequency distributions based on velocity, computer simulation was performed using the MC method. Computational results are shown in Figure 5. The power-law exponent in the vertical direction was 1.43 for females (least mean square = 0.99) and 1.17 for males (0.99) and that in the horizontal direction was 1.02 for females (0.98) and 1.89 for males (0.99). The male-to-female ratio of the power-law exponent was 0.82 in the vertical direction and 1.85 in the horizontal direction.

**Figure 5.**
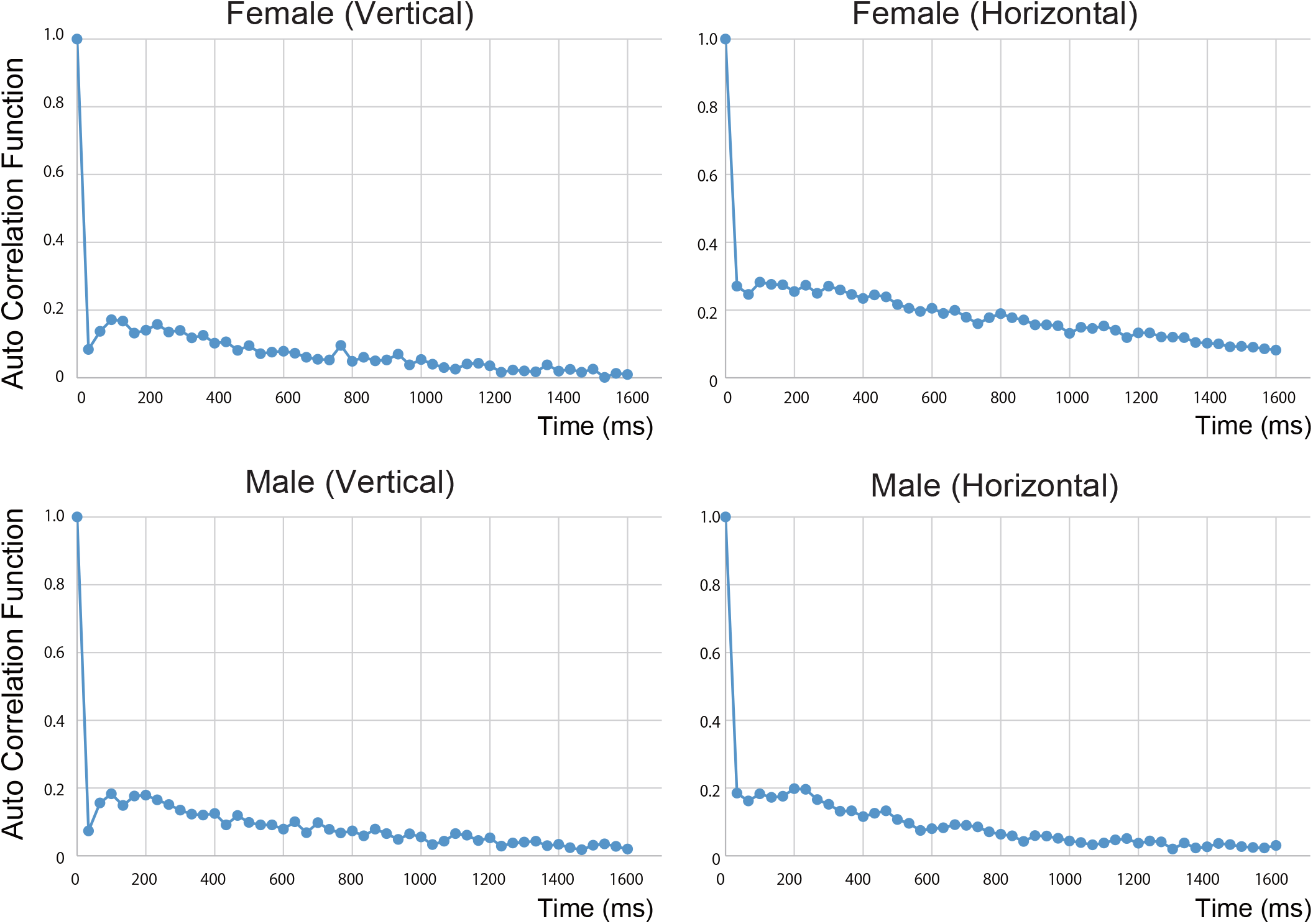
Mean squared displacement of model MC1. The mean squared displacements (MSDs) of females (left) and males (right) of model MC1 are plotted along the vertical (top) and horizontal (bottom) axes (logarithmic scales, *n* = 28). Best-fit lines using a power-law approximation are shown in black. The power-law exponent (α) and least mean square (*R*^2^) are shown in the graphs. The data shown represent the means ± SEM.

### Frequency of move-step length and MC2

The frequency distribution of the swimming speed (move-step length, mm/video frame; n = 7,168 for each sex) is shown in Figure 6. The results clearly illustrate a high frequency in the lower move-step length, which gradually decreased as the move-step length increased. In both female and male individuals, the frequency distributions were well fitted by exponential functions (Figure 6, right). The slope of the regression lines for the vertical direction was −5.03 for females (least mean square = 0.97) and −6.08 for males (0.99). The slope of the regression lines for the horizontal direction was −4.55 for females (0.98) and −5.59 for males (0.96). For both directions, the slope of the regression line was greater for females than for males. Significant differences in the mean move-step lengths in each direction were observed between females and males (females and males in the vertical direction: 0.26 ± 0.0027 mm/video frame and 0.21 ± 0.0022 mm/video frame, respectively; and females and males in the horizontal direction: 0.28 ± 0.0030 mm/video frame and 0.25 ± 0.0027 mm/video frame, respectively; *p* < 0.05, unpaired *t*-test). In both directions, the mean move-step lengths of females were longer than those of males.

**Figure 6.**
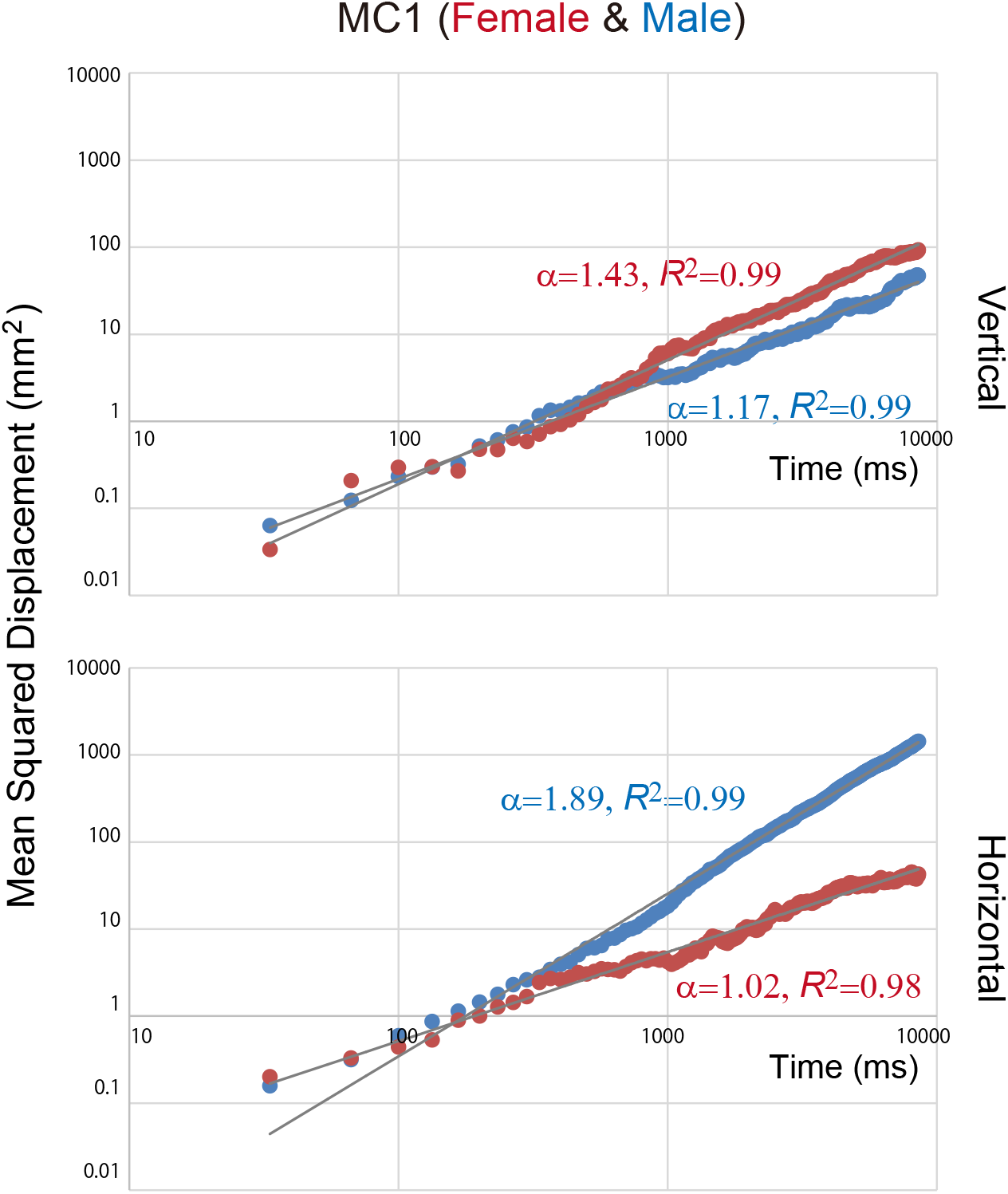
Frequency distribution of move-step length. The move-step lengths derived from 7,168 velocities are plotted in linear (left) and single logarithmic (right, log-linear model) scale plots. The data for females (red) are superimposed on the data for males (blue). Bin width, 0.12 mm. Best-fit lines using an exponent function are shown in black. The exponent function and least mean square are shown in the plots.

MC simulations using model 2 (MC2), which was derived from swimming speed, were conducted. Plus or minus signs were randomly assigned to randomly selected move-step lengths, the re-signed values were used as a data set for swimming velocity and MSDs were calculated. The power-law exponent in the vertical direction was 0.92 for females (least mean square = 0.98) and 0.92 for males (0.90). The power-law exponent in the horizontal direction was 1.07 for females (0.97) and 1.13 for males (0.97). The male-to-female ratio of the power-law exponent was 1.00 in the vertical direction and 1.06 in the horizontal direction.

### Size of a cluster with an identical sign and MC3

In addition to regular jumping motions, *D. magna* occasionally makes a large change in the direction of movement (left/right or up/down) via turning behaviour. These turning phenomena can be represented by the direction of movement (left/right or up/down). We accordingly calculated the sign of the direction of movement. Dominant and recessive directions were defined for each data set, with the dominant direction represented by a plus sign (Figure 7A, see Methods for details). Although the turning behaviours have a large effect on the global trajectory of *D. magna,* capturing the turning motion from local trajectories based on the time unit of the video frame is difficult. For this reason, to visualise the turning motion, we used a parameter termed SCIS, (Figure 7A, see Methods for details). The frequency distribution of the parameter SCIS is shown in Figure 7B. A long-tailed distribution (arrow) was found for positive-sign movements of males. In Figure 7C, the dominant area of the frequency distribution is shown on logarithmic scales. The power-law exponent of the frequency distribution in the vertical direction was −3.15 for females (least mean square = 0.98) and −3.12 for males (0.98). The power-law exponent in the horizontal direction was −3.22 for females (0.96) and −1.87 for males (0.99). The mean distance in the horizontal direction was 118.84 ± 18.32 mm for males, significantly greater than that for females (25.63 ± 4.54 mm). In contrast, the mean length of the movements of males in the vertical direction was 29.84 ± 5.41 mm, not significantly different from that of females (22.92 ± 2.63 mm).

**Figure 7.**
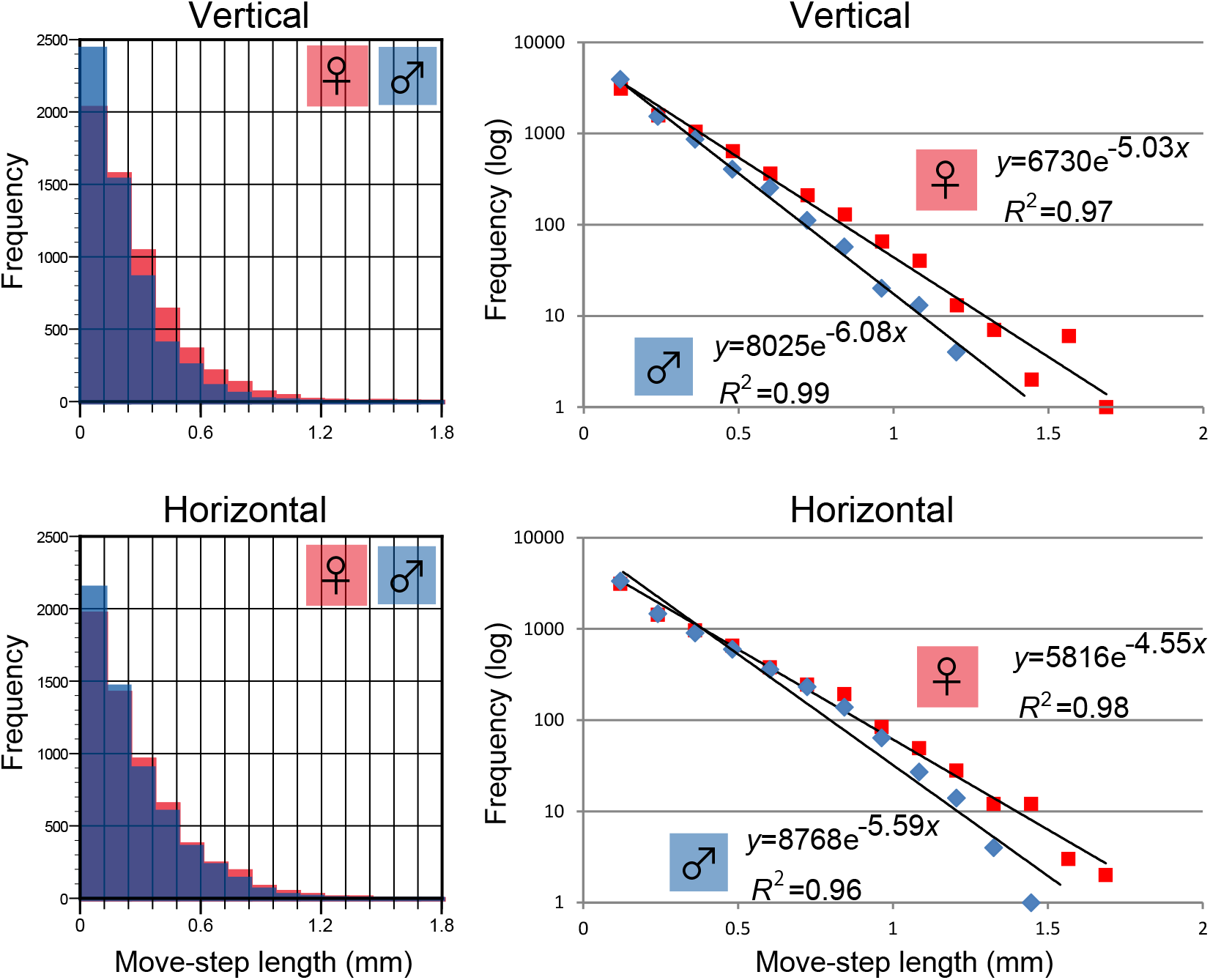
SCIS analysis. (A) Brief overview of the definition of the dominant and recessive directions and durations with an identical sign. The major direction is defined as dominant and is indicated by the plus sign (refer to Methods for details). (B) The frequency of the duration with an identical sign derived from 7,168 velocities is plotted at the bottom of the graphs. Frequencies greater than 1,500 ms and less than −1,500 ms were added to the respective end of the bin. Bin width, 166.5 ms. The dominant area is shown in dark grey, and the recessive area is shown in pale grey. In the insets, the tail regions of graphs are enlarged. (C) The dominant area of B is shown on logarithmic scale. Female *D. magna* is shown in red and male *D. magna* in blue. Best-fit lines using a power-law approximation are shown in black. The power-law exponent (α) and least mean square (*R*^2^) are shown in the plots. .

Assuming an MC in reference to the SCIS frequency distributions, MC simulations using model 3 (MC3), which was derived from SCIS, were performed to calculate MSDs (Figure 8). The power-law exponent of the MSD in the vertical direction was 1.79 for females (least mean square = 0.98) and 1.82 for males (0.99). The power-law exponent in the horizontal direction was 1.38 for females (0.98) and 1.96 for males (0.99). The male-to-female ratio of the power-law exponent was 1.02 and 1.42 in the vertical and horizontal directions, respectively. Characteristics of the two time scales observed in the motions of females in the horizontal direction (Figure 3, bottom, red) appear to be reproduced by the MC3 model. The power-law exponents of the MSDs of *D. magna* and three MC models are summarised in Table 1.

**Table 1.** Summary of three MC models. The power-law exponents of MSDs of *D. magna* and three MC models are summarised. V: vertical direction, H: horizontal direction, M/F: ratio of male divided by female.

|  |  | *D. magna* | MC1 | MC2 | MC3 |
| --- | --- | --- | --- | --- | --- |
| **Female** | **V** | 1.58 | 1.43 | 0.92 | 1.79 |
|  | **H** | 1.29 | 1.02 | 1.07 | 1.38 |
| **Male** | **V** | 1.39 | 1.17 | 0.92 | 1.82 |
|  | **H** | 1.82 | 1.89 | 1.13 | 1.96 |
| **M/F** | **V** | 0.88 | 0.82 | 1.00 | 1.02 |
|  | **H** | 1.41 | 1.85 | 1.06 | 1.42 |

**Figure 8.**
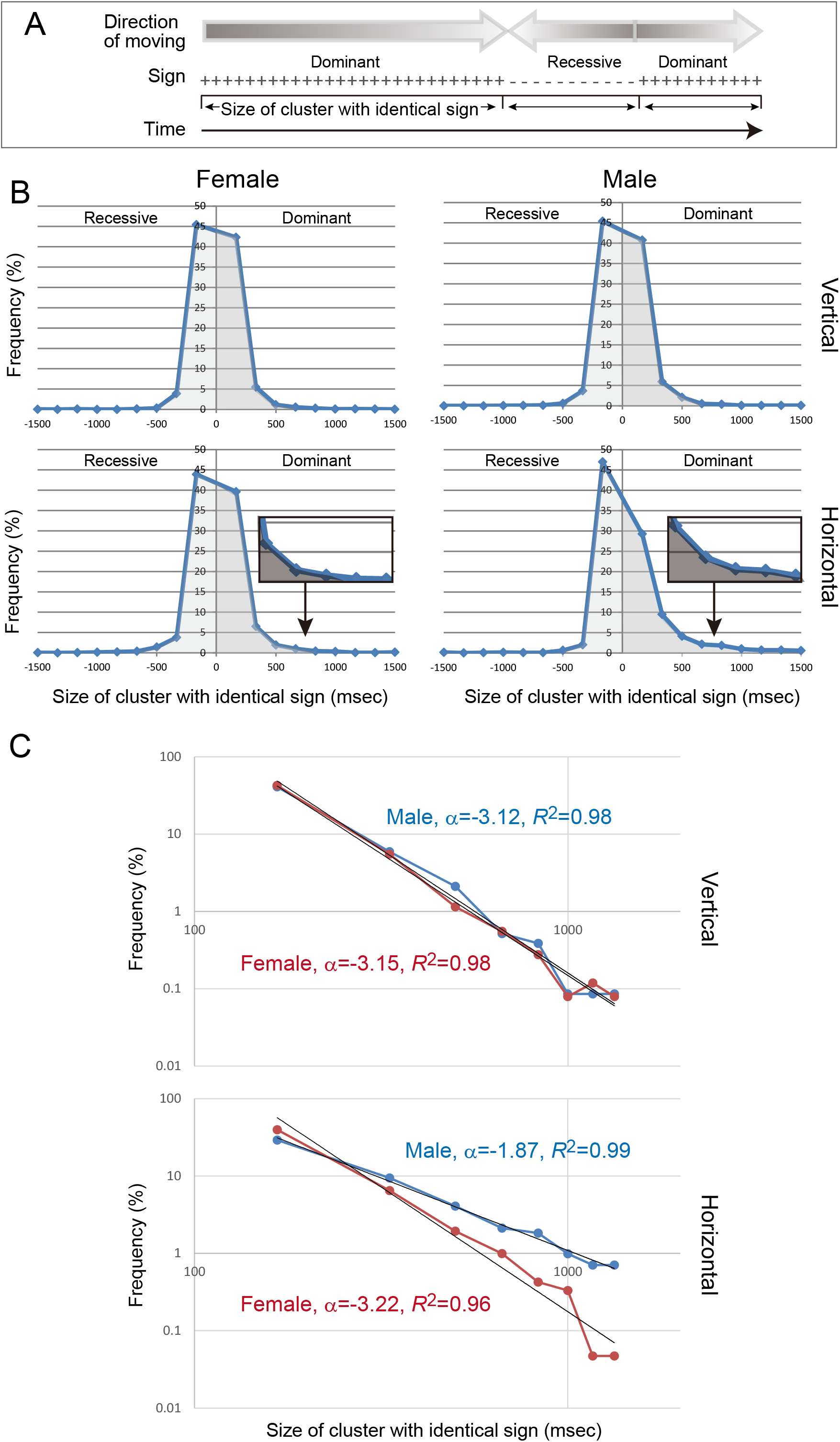
Mean squared displacement of model MC3. The mean squared displacements for females (red) and males (blue) based on model MC3, which was derived from the duration with an identical sign, are plotted along the vertical (top) and horizontal (bottom) axes (logarithmic scales, n = 28). A constant arbitrary speed (7.2 mm/s) was used for this simulation. Best-fit lines using a power-law approximation are shown in black. The power-law exponent (α) and least mean square (*R*^2^) are shown in the plots.

### Net-to-gross displacement ratios

NGDRs (refer to Figure 9A) were calculated from three data sets: (1) the raw data derived from *D. magna*, (2) MC1 and (3) MC3 (Figure 9B). NGDRs provide a measure of the relative linearity of the swimming path of the plankton; lower NGDRs imply more curved trajectories.

**Figure 9.**
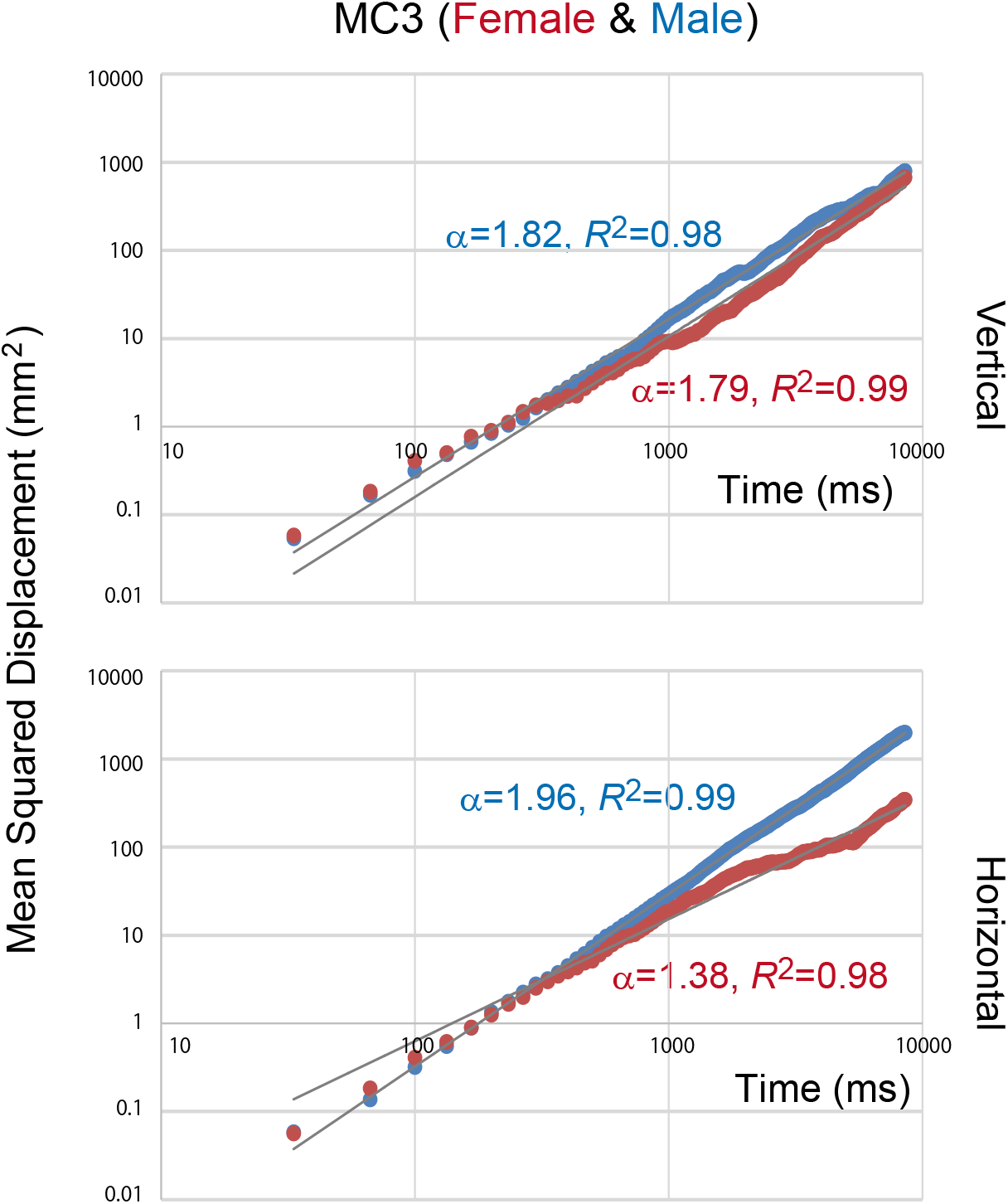
Net-to-gross displacement ratios of *D. magna* and two MC models. (A) Brief overview of the definition of the net-to-gross displacement ratios (NGDRs). Net displacement (ND) is the straight line distance between the initial and final locations, and gross displacement (GD) is the sum of the distance D*i*. (B) NGDRs observed in the raw data derived from *D. magna* are shown relative to those from MC1 and MC3. The term ‘fv’ indicates the vertical direction of females, ‘fh’ indicates the horizontal direction of females, ‘mv’ indicates the vertical direction of males and ‘mh’ indicates the horizontal direction of males.

The NGDRs of the raw data from females were 0.32 ± 0.027 in the vertical and 0.32 ± 0.050 in the horizontal direction. The NGDRs of the raw data from males were 0.33 ± 0.041 and 0.70 ± 0.053 in the vertical and horizontal directions, respectively. The male-to-female ratios of the NGDRs were 1.03 in the vertical and 2.19 in the horizontal direction. These results indicate that NGDR in the horizontal direction for males was significantly higher than the other NGDRs (*p* < 0.01, unpaired *t*-test). Thus, the linearity of the swimming path of males in the horizontal direction was significantly greater than that of the other direction and sex.

This specificity was reproduced in the two MC models. The NGDRs of MC1 for females were 0.13 ± 0.015 and 0.078 ± 0.0087 in the vertical and horizontal directions, respectively, and those for males were 0.11 ± 0.011 and 0.54 ± 0.013, respectively. The male-to-female ratio of the NGDR was 0.85 in the vertical and 6.92 in the horizontal direction. The NGDRs of MC3 for females were 0.31 ± 0.029 and 0.28 ± 0.030 in the vertical and horizontal directions, respectively, and those of males were 0.36 ± 0.025 and 0.77 ± 0.025, respectively. The male-to-female ratios of the NGDR were 1.16 in the vertical and 2.75 in the horizontal direction. These results indicate that based on the MC models, the NGDR in the horizontal direction for males was distinctly higher than the other NGDRs (*p* < 0.01, unpaired *t*-test).

## Discussion

The motions of female and male *D. magna* individuals were tracked in two-dimensional coordinates; then, three diffusion parameters (kernel density estimates, MSDs and NGDRs) were calculated. All of these parameters showed that male adult *D. magna* individuals exhibit greater diffusiveness in the horizontal direction than females. This male-dominant diffusion pattern was computationally reproduced by the MC method in which the frequency map of the velocity was considered as a probability distribution. Then, the factors that might have contributed to the particularly high diffusiveness of males in random motions were analysed using the MC method. The frequency maps of the migration length per unit time (absolute value of the moving velocity) were nearly equal between females and males. The frequency distribution of the swimming speed in females and males did not explain the higher diffusion of males in the horizontal direction. Because these organisms often make drastic changes in the direction of movement, the frequency of turning behaviour (SCIS) was calculated. When the frequency maps of SCIS were considered as a probability distribution, the MC computational simulations reproduced the observed sexual difference in diffusiveness, showing that male *D. magna* likely predominantly diffuse in the horizontal direction by changing the frequency of turning behaviour. Regardless of the low autocorrelation found for velocity, the sexual difference was reproduced in model MC3, which is a type of persistent random walk. This result appears to show inconsistency. However, variation of move-step length was not included in MC3 (move-step length was constant in MC3). Considerable randomness and variation of move-step length may have contributed to the low autocorrelation. Furthermore, two-valued directions (plus or minus) in SCIS possibly enhance the persistency. In either case, it should be noted that SCIS analysis extracts the persistency from low-autocorrelation data. For detailed evaluation of this question, comparison with other analyses using data with higher spatial and temporal resolution or data to which a smoothing method applies will be helpful (Hen et al., 2004; Postlethwaite and Dennis, 2013).

The three abovementioned indices of the autonomous free diffusion of *D. magna* describe only a portion of their broad spectrum of ecological dynamics and were calculated using distinct processes. Nonetheless, all of these indices commonly captured the greater diffusiveness of males in the horizontal direction. Such preferred movement by males in the horizontal direction has been observed in another daphnid species (*D. pulicaria*) (Brewer, 1998). Females exhibited a strong vertical swimming component, ranging from −60 to +75 from the horizontal, whereas male swimming behaviour was primarily linear and horizontally oriented, with swimming angles deviating no more than ±15 from the horizontal. Furthermore, very different motility was observed between female and male copepod crustaceans (*Centropages typicus* and *Pseudocalanus elongates*) (Kiørboe and Bagøien, 2005). Males were more directionally persistent than females based on root-mean-square net distance. Thus, the comparatively high diffusion of males in the horizontal direction may be a general phenomenon in zooplankton. Over 200 species of daphnids (Kotov et al., 2012) and over 13,000 species of copepods (Boxshall and Defaye, 2008) have been discovered to date; however, there are few zooplankton in which motion patterns have been investigated. Further verification of their behaviour is required.

### Random motion of D. magna

In this study, we investigated the autonomous movement pattern of *D. magna* individuals under uniform conditions with no water stream, a constant light cycle and temperature. To avoid chemical and physical influences from animals of the opposite sex, the individual motion of daphnids was measured while they were swimming together with those of the same sex. The autonomous movement pattern of each zooplankton in such a uniform environment in the laboratory was measured by various researchers. This characteristic was detected and expressed using various mathematical models (Okubo and Levin, 2001; Seuront and Strutton, 2003), which are useful for the analysis and comparison of the patterns of animal movement in any species.

In this study, MC simulation analysis was performed using speed distribution data from raw tracking data considered as a probability distribution. As a result, the differences between males and females mentioned above were reproduced. Thus, it can be inferred that it is appropriate to analyse as a Markov process the difference in diffusion between males and females. Reproducibility using the MC model indicates that only the present position is used as past information determining the future motion of plankton and that the future motion follows specific probability distributions. In our recent study, when visual stimulation was generated via a MC method in which speed distribution data from raw tracking data of female *D. magna* were considered as a probability distribution, visual stimuli promoted the feeding behaviour of medaka fish (*Oryzias latipes*) (Matsunaga and Watanabe, 2012). This finding suggests that the Markov process observed in the movement of *D. magna* is an important biological phenomenon. Furthermore, in this study, movement of *D. magna* was separated into migration distance per unit time and frequency of turning behaviour. The frequency of turning qualitatively explained the behaviour of males.

Turning behaviour has been observed in small organisms such as *Caenorhabditis elegans* (Ohkubo et al., 2010; Srivastava et al., 2009), *Paramecium* (Nakaoka et al., 2009) and *Drosophila* (Censi et al., 2013). These turning behaviours occur even under spatially and temporally uniform conditions, indicating that these are self-motivated and voluntary behaviours. Theoretical studies suggest that stochastic behaviours are important for efficiently searching for prey and mates (Bartumeus et al., 2008). A similar turning phenomenon was observed in feeding female *D. magna*, phytoplankton distributed throughout the aquatic systems of lakes and ponds. The male *D. magna* could have adapted to the rapidly changing habitat environment by slightly reducing their probability of turning. To clarify the biological meaning of this stochastic process, the presence of some internal mechanism regulating the probability distribution of turning behaviour is suggested.

Sex-specific morphological characteristics might be associated with the generation of sex-specific behavioural differences. Indeed, mature male daphnids are distinguishable from females not only by their testes and sperm but also by several apparently distinctive morphological characteristics, such as a smaller body size, elongated first antennae and a copulatory hook at the first thoracic leg (Brewer, 1998; Mitchell, 2001). However, the causal relationships of the sexual differences between behavioural and morphological characteristics in daphnid species remain largely unknown.

### Acquisition of genetic heterogeneity

As mentioned above, the higher diffusiveness of male zooplankton has been observed in multiple species, and the underlying dynamic regulation of turning has been reported in various animal species (Censi et al., 2013; Nakaoka et al., 2009; Srivastava et al., 2009). Thus, discussing the biological significance of this difference in diffusiveness would be useful, even though a conclusion on this issue is beyond the scope of the present study. Here we propose simply that the high diffusiveness of male *D. magna* is closely related to ensuring genotype heterogeneity.

As shown in Figure 2C, female *D. magna* exhibit heavier recursiveness to the point of origin, indicating that the motion of females exhibits less diffusion than that of males. Under normal conditions, female *D. magna* generate many offspring that are genetic clones of the mother. Because *D. magna* have no sex chromosome, both female and male offspring are genetically identical to the mother. Given that male *D. magna* individuals exhibit diffusion similar to that of females, males are more likely to encounter clones of their own family. We accordingly hypothesise that the comparatively high diffusive motion of male *D. magna* leads to displacement far from their generation, thus impeding contact with its family members. As a result, they can produce resting eggs that exhibit high genetic variation. It appears reasonable that water fleas living in shallow ponds concentrate their energy in horizontal directions.

Traditionally, behavioural patterns of daphnids have been demonstrated by population (mass culture) levels under laboratory and/or natural environments (Beaver et al., 2018; Dodson, 1988). Recent technologies have enabled the individual-level behavioural analysis by tracking video system, and it can be applied for the toxicological and risk assessment of chemicals as a novel endpoint (e.g., swimming velocity and trajectory) in addition to, for examples, survival rate and fecundity of *D. magna* (Bownik et al., 2019; 2020; Felice et al., 2019; Noss e al., 2013). Although *D. magna* has been used in ecotoxicological studies as a representative surveillance organism to monitor changes in water quality (Martins et al., 2007), little is known about the basic mechanisms involved in its behaviour. In this study, we found that male individuals travelled in a diffuse horizontal direction relative to females. Furthermore, this male-specific diffusion pattern was reproduced by mathematical modelling using MC simulation, in which a frequency map of their turning behaviour was used as a probability distribution. Although it is necessary to clarify strain differences of behavioural patterns (Oda et al., 2007), current data provides an insight of new endpoints to OECD Test Guidelines no. 211 using daphnids (OECD, 2012), enabling the validation of chemical toxicity based on behaviour. Likewise, we recently found useful *Daphnia* strains that can produce female or male in response to day-length differences in the *D. pulex* (Toyota et al., 2015) and *D. magna* (Toyota et al., 2019; 2021). These strains will be used for comparative behavioural analysis of inter- and intra-species between sexes, and moreover 3D tracking approaches (Bianco et al., 2013) will shed light on the well-conserved innate behavioural patterns of daphnids. Our findings provide the first insight into the behavioural characteristics of female and male adult *D. magna* individuals and lay the foundation for advancing our understanding of the dynamics of daphnid populations and the toxic effect of chemicals released into the aquatic environment.

## Materials and Methods

### Daphnia magna strain and rearing conditions

A *D. magna* strain (Belgium clone) was obtained from the National Institute for Environmental Studies (NIES; Tsukuba, Japan) (Oda et al., 2006). Stock populations were maintained as reported previously (Tatarazako et al., 2003). Briefly, they were housed in a 30 L aquarium (containing 20 L of housing water) and maintained in an incubator at 24 ± 1 °C on a 14/10 h light/dark cycle (with lights on from 08:00 to 22:00). The housing water was prepared by mixing artificial sea salt into deionised H_2_O (18 g/60 L; Tetra Marine Salt Pro, Tetra Japan, Tokyo, Japan) (Matsunaga and Watanabe, 2012). A chlorella (*Chlorella vulgaris*) suspension (0.3 mL/10 L; Chlorella Industry, Tokyo, Japan) was added to the housing tank as a food source once a day at 08:30. This strain occasionally produces male offspring in addition to female offspring under overpopulation conditions. Male juveniles were easily distinguished based on their elongated first antennae (Olmstead and LeBlanc, 2000). Two- to three-week-old male and female animals were used in subsequent experiments.

### Motion recording

Cuboid clear plastic aquaria [interior dimensions: 9.5 cm (width) × 1.0 cm (length) × 9.5 cm (vertical depth)] were used as test tanks for the motion analysis of *D. magna*. The aquaria were filled with housing water to a vertical depth of 8.0 cm. The temperature of the tank water was maintained at 25 ± 2 °C by air conditioning in the experiment room. The bottom of each tank was covered with a black plastic mat to prevent light reflection. The sides of the tanks, excluding the side on which the digital video camera (Himawari GE60; Library, Tokyo, Japan) was positioned, were covered with black rubber to prevent excessive illumination and light reflection. The test tank was placed in a dark room. Illumination at the surface of the water was adjusted to 3,000 lx using three white fluorescent lamps placed outside the tank above the surface, excluding the side on which the video camera was positioned. Male or female (*n* = 10–20 each) *D. magna* individuals were transferred to the tank, and motion was recorded from the side of the tank using the digital video camera. The experiments on each female and male were repeated nine times (18 trials in total). Video images (640 × 480 pixels) were recorded at 30 frames per second (fps) and were quantified with DIPP-Motion 2D motion analysis software (DITECT, Tokyo, Japan). The coordinates of the centre of mass of the individuals were automatically tracked in each video frame. The coordinates were obtained as integer numbers (horizontal coordinates from 0 to 639; vertical coordinates from 0 to 479). Three or four *D. magna* individuals were tracked in each experiment. Data points in which individuals collided against each other or the tank walls were excluded. Thus, sexual differences in swimming behaviours were studied in reference to the autonomous movement pattern of each individual in a uniform environment. Consecutive coordinates (257) for each daphnid were treated as a data set, and 28 data sets were collected for each sex. The frequency distribution of the velocities showed a single peak with wide tails located at either side (Supplemental Figure 1). The maximum values were 13 pixels (horizontal) and 13 pixels (vertical) for females and 9 pixels (horizontal) and 10 pixels (vertical) for males. The minimal values were −14 pixels (horizontal) and −14 pixels (vertical) for females and −12 pixels (horizontal) and −9 pixels (vertical) for males. In the following analysis, these integral data were used and then finally converted to millimetre scale.

### Kernel density estimation

In total, 7,196 coordinates (240 sets, 257 consecutive coordinates per individual *D. magna*, *n* = 28 for each sex) of the calculated centre of mass of *D. magna* were obtained. The consecutive coordinates were normalised to the given halfway point (the 129^th^ coordinate), which was set as vertical coordinate 50 and horizontal coordinate 50 (Figure 1, top). Nonparametric kernel density estimation (Silverman, 1986; Fortmann-Roe et al., 2012; Worton, 2002) of the normalised trajectory data was performed using a quartic kernel function with a bandwidth of 12 mm. The kernel density estimator at coordinate **x**is given as follows:

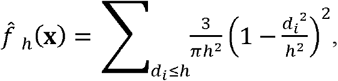

where *h* is a smoothing parameter termed bandwidth and *d*_*i*_ is the distance from coordinate **x** to coordinate **x***i*. The kernel density estimator was applied to 12,100 (110 vertical × 110 horizontal) coordinates and the two-dimensional density map was visualised with terra colours using R software (http://www.r-project.org/).

### Motion analysis

The movement distance per frame was calculated from a series of coordinates and converted to swimming velocity (mm/s), which was calculated separately for the horizontal axis (with increasing values to the right) and the vertical axis (with increasing values upward). The swimming speed was calculated as the absolute value of the swimming velocity. Mean squared displacement (MSD) was calculated from a series of coordinates as follows:

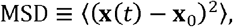

where **x**_0_ is the initial position of a plankter and **x**(*t*) is the position of the individual at a specific time *t*.

The discrete autocorrelation Rs at lag *n* were computed as follows:

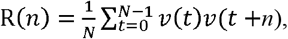

where *v*(*t*) is the velocity of an individual plankter at a specific time *t*. The normalised autocorrelation function (ACF) was calculated as follows:

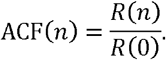

Net-to-gross displacement ratios (NGDRs) (Figure 9) were computed as follows (Busley, 1984):

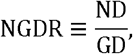

where ND (mm) and GD (mm) are the net and gross displacements, respectively, of an individual; these measures correspond to the shortest distance between the starting and ending points of the trajectory and the actual distance travelled by an individual, respectively. NGDRs were computed at the finest available resolution (1/30 s) for each track. The statistical significance of the results of NGDR was determined using unpaired (two-tailed) *t*-tests. The threshold for statistical significance was *p* < 0.05. Values are reported as means ± SEM.

One-dimensional velocities in each set of 255 data have either plus or minus signs. The most frequent sign was defined as the dominant direction. Dominant and recessive directions were determined for each set of data, and the dominant direction was represented anew by a plus sign. The movement distance per frame was converted to values accompanied by signs referring to the dominant and recessive directions. The size of a cluster with an identical sign (SCIS) was calculated as the duration for which identical signs were continuous (Figure 7A). Periods of zero speed were assigned the sign of the preceding period.

### Markov chain simulation

Computer simulations using the MC method were performed. The virtual zooplankton models were assumed to be uncorrelated random walks referencing the motion parameters of *D. magna*. First, independent random variables, *v*_1_, *v*_2_, *v*_3_,…, *v*_255_, were taken with each variable generated by a randomiser referencing the discrete probability distributions of velocities (model MC1), move-step lengths (model MC2) or SCISs (model MC3). Then, we defined

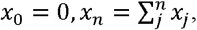

where *x*_0_ is the initial position of a model plankter and *x*_*n*_ is the position of the individual at a discrete specific time j. Finally, the MSDs and NGDRs of each set of progressions {*x*_*n*_} were calculated. Progressions {*v*_1_, *v*_2_, *v*_3_,…, *v*_255_} for each MC model were generated as follows.

#### MC1

1) The raw velocity data from 28 *D. magna* individuals were collected into a data set (255 velocities per individual; *n* = 7,140 in total for each direction). 2) In total, 255 values were randomly selected from each data set. The values were selected with repetitions. 3) A random selection of values was repeated 28 times for each data set. The reselection of an identical data point was not excluded.

#### MC2

1) The raw move-step length data from 28 *D. magna* individuals were collected into a data set. 2) In total, 255 values were randomly selected from each data set. 3) A random selection of values was repeated 28 times for each data set. Values were selected with repetitions. 4) A plus or minus sign was randomly assigned to the selected move-step length.

#### MC3

1) The raw SCIS cluster data from 28 D. magna individuals were collected into a data set. Each cluster included a sign (plus or minus) and a discrete size. 2) A cluster was randomly selected from each data set. 3) The selected cluster was converted to a progression where 1 (if the cluster sign was plus) or −1 (if the cluster sign was minus) was sequenced by the length of the cluster size. 4) Steps 2 and 3 were performed again, and the first and second progressions were combined. Clusters were selected with repetitions. 5) Steps 2–4 were repeated until the length of the progression reached 255. 6) Members of the progression were multiplied by an arbitrary constant value (2 pixels/video frame = 7.2 mm/s). 7) The generation of the progressions was repeated 28 times for each data set.

## Acknowledgements

We are grateful to Drs. Hitoshi Miyakawa (Utsunomiya Univ.) and Chizue Hiruta (Hokkaido Univ.) for their helpful comments on this manuscript, to Dr. Kei-ichi Okunuki (Nagoya Univ.) for helpful advice on the GIS method, to Dr. Hiroshi Koyama for helpful advice on the random walk models, and to Ms. Mie Watanabe for her technical assistance in preparing the manuscript.

## Competing interests

The authors declare no conflicts of interest.

## Funding

The Ministry of Education, Culture, Sports, Science, and Technology of Japan supported this work.

---

**Supplemental Figure 1. Frequency distribution of velocity.**

A frequency histogram of the velocities derived from 7,168 vectors derived from female or male zooplankton. The velocity was calculated separately in the horizontal and vertical axes. Bin width = 1 pixel/video frame (0.12 mm/33.3 ms).

**Supplemental Movie 1. Female *D. magna*.**

An example of a female *D. magna*. Width of the movie is 640 pixels (76.8 mm), and height is 480 pixels (57.6 mm). Frame rate is 30 fps.

**Supplemental Movie 2. Male *D. magna*.**

An example of a male *D. magna*. Width of the movie is 640 pixels (76.8 mm), and height is 480 pixels (57.6 mm). Frame rate is 30 fps.

